# Re: Concerns Regarding the Validation of ZYS-1 as a *Bona Fide* ADAR1 Inhibitor

**DOI:** 10.1101/2025.03.07.641892

**Authors:** Min Zhang, Xiangyu Deng, Yang Gao, Jin Wang

## Abstract

Wang et al. in *Nature Cancer* report ZYS-1 as an ADAR1 inhibitor and present biochemical and cellular evidence to support ZYS-1’s direct interaction with and inhibition of ADAR1, including enzymatic assays, surface plasmon resonance (SPR), drug affinity responsive target stability (DARTS), and cellular thermal shift assays (CETSA). We raise critical methodological concerns regarding the validation of ZYS-1 as a *bona fide* ADAR1 inhibitor. Our primary concern relates to the misuse of an adenosine deaminase (ADA) activity assay kit for ADAR1 enzymatic activity characterization. Using established biochemical assays with direct inosine quantification, we evaluated ZYS-1’s inhibitory activity against ADAR1 using physiologically relevant RNA substrates. The compound showed no inhibitory activity at concentrations up to 1 mM. We demonstrate that the ADA assay kit used by Wang et al. does not reflect ADAR1-specific activity, and ZYS-1 is unable to inhibit ADAR1 in well-validated biochemical assays. We urge for further validation using ADAR1-specific assays to establish ZYS-1 as a genuine ADAR1 inhibitor.

## Main Text

Dear Editor,

We read with great interest the recent article by Wang et al. in *Nature Cancer* reporting the discovery of ZYS-1 ^1^, described as a potent small-molecule inhibitor of Adenosine Deaminase Acting on RNA 1 (ADAR1).

ADAR1 is an RNA-editing enzyme critical for the post-transcriptional deamination of adenosine to inosine in double-stranded RNA (dsRNA) ^2^. It exists in two isoforms: constitutively expressed p110, localized to the nucleus, and interferon-inducible p150, which mainly resides in the cytoplasm ^3^. The p150 isoform plays a key role in dampening innate immune responses by editing immunogenic dsRNA, thereby preventing aberrant activation of cytosolic RNA sensors. The landmark study from the Hanning group reported that loss of ADAR1 in tumors overcomes resistance to immune checkpoint blockade ^4^, which has garnered significant interest in both academia and industry to develop ADAR1 inhibitors or degraders ^5^. Despite this interest, pharmacological modulation of ADAR1 remains elusive, with some misconceptions in the field—for instance, a study incorrectly classified spliceosome inhibitors as ADAR1 inhibitors ^6^.

Wang et al. report ZYS-1 as an ADAR1 inhibitor and present extensive biochemical and cellular evidence to support ZYS-1’s direct interaction with and inhibition of ADAR1, including enzymatic assays, surface plasmon resonance (SPR), drug affinity responsive target stability (DARTS), and cellular thermal shift assays (CETSA) ^1^. While these findings are intriguing, we wish to highlight critical methodological concerns that call into question the validity of ZYS-1 as a bona fide ADAR1 inhibitor.

Our primary concern of Wang et al. in *Nature Cancer* ^1^ relates to the misuse of the adenosine deaminase (ADA) activity assay kit for ADAR1 enzymatic activity characterization, which is the foundational experiment to demonstrate ZYS-1 as a *bona fide* ADAR1 inhibitor. The enzymatic inhibition assay in their Figure 5b employs a commercial ADA activity kit to measure ADAR1 activity. However, ADA and ADAR1 are functionally distinct enzymes: ADA catalyzes the deamination of free adenosine to inosine ^7^, whereas ADAR1 edits adenosine within dsRNA substrates ^2^.

Although the mechanism of the Abcam ADA assay kit (Abcam ab204695) used in Wang et al. ^1^ was not disclosed, similar assay kits revealed their assay principle ^8^. Based on the assay components in the Abcam kit, we rationalize that the kit utilizes a coupled enzyme system, in which free inosine is converted to hypoxanthine by purine nucleoside phosphorylase (PNP) ^9^. Xanthine oxidase (XOD) then converts hypoxanthine into xanthine by releasing H_O_ ^10^, which can be quantified by the OxiRed probe through fluorogenic signals ^11^. Importantly, PNP does not convert inosine within dsRNA into hypoxanthine ^9^, making this assay fundamentally unsuitable for assessing ADAR1’s RNA-editing activity. We contacted Abcam technical support, who confirmed that this kit is not designed or validated for ADAR1 activity measurements.

We noticed that in the preprint published by the same authors in 2021 ^12^, the ADA assay kit was directly used to measure ADAR1 enzymatic activity, suggesting a confusion about ADA and ADAR1 as two distinct enzymes. In the current published study ^1^, human Glioma-associated oncogene 1 (GLI1) mRNA was used as the substrate instead of the ADA substrate from the kit to determine ADAR1 and ADAR2 deaminase activities. However, the authors fell short of using the edited product of inosine-containing GLI1 RNA to establish a standard curve. Additionally, although p110 and p150 are two distinct forms of ADAR1, the authors did not specify which form was used in the biochemical assay. It is also unclear whether the purchased recombinant ADAR1 is functionally active.

To independently test whether the ADA assay kit could detect ADAR1 editing activity, we incubated a series of concentrations of inosine-containing dsRNA (GLI-dsRNA-Product, the editing product of ADAR1, **Figure 1A**) and inosine-non-containing dsRNA (GLI-dsRNA-2, the editing substrate of ADAR1, **Figure 1A**) with ADA converter, converter enzyme VIII, and the OxiRed probe. We observed only background level fluorescence signals, while free inosine generated a linear standard curve as expected (**Figure 1B**). This experiment demonstrates that the ADA kit can quantify free inosine but is unable to respond to inosine inside dsRNA, making it unsuitable for use as an activity assay for ADARs.

**Figure 1.**
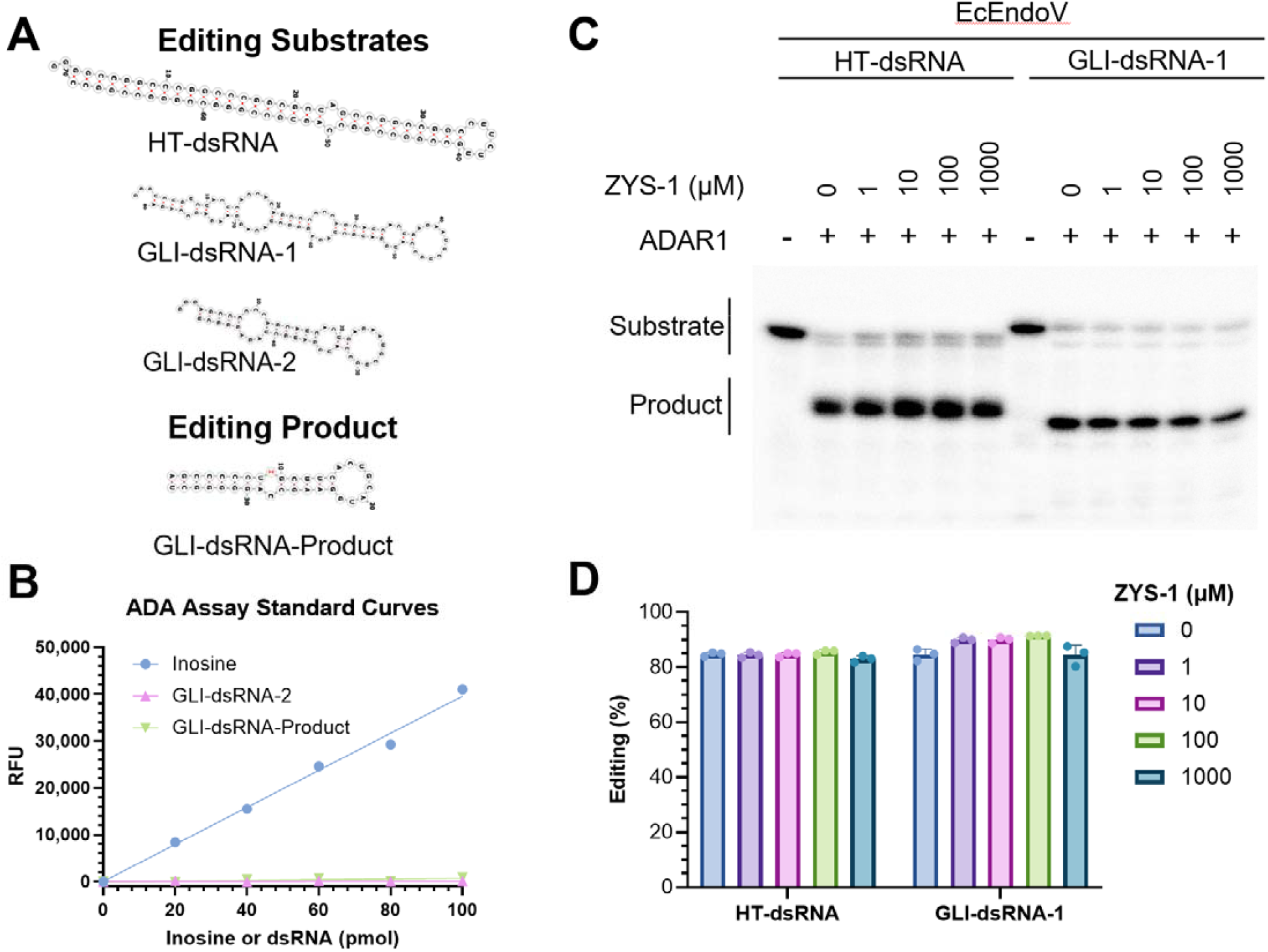
Evaluation of ZYS-1 as a *bona fide* ADAR1 inhibitor. (A) The sequences and secondary structures of dsRNAs used in our assays. HT-dsRNA, GLI-dsRNA-1, GLI-dsRNA-2 are the ADAR1 editing substrates, containing adenosine editing sites. GLI-dsRNA-Product is the ADAR1 editing product, containing inosine at the editing site. (B) Responses of adenosine deaminase activity (ADA) assay kit to inosine, ADAR1 editing substrate GLI-dsRNA-2 and ADAR1 editing product GLI-dsRNA-Product. (C) Representative gel images showing the RNA editing activity of human ADAR1 in the presence or absence of the ZYS-1 compound. Reactions were performed with 50 nM HT-dsRNA or GLI-dsRNA-1 and 150 nM ADAR1, with ZYS-1 concentrations as indicated. The observed product results from ADAR1-mediated RNA editing, followed by cleavage with EcEndoV, which specifically targets inosine-containing RNA. (D) Quantification of normalized RNA editing activity from (C). Each experiment was independently repeated at least three times.

We recently established the expression of a ADAR1 p150 variant and developed a dsRNA editing functional assay ^13^. In this assay, the editing product (inosine-containing dsRNA) is cleaved by *Escherichia coli* Endonuclease V (EcEndoV), followed by gel electrophoresis to quantify the ratio between the fragments and full-length dsRNA. Using this approach, we found that ADAR1 editing is both sequence and RNA duplex length-dependent but can tolerate mismatches near the editing site. Additionally, we have solved high-resolution structures of ADAR1-RNA complexes which, coupled with mutagenesis studies, revealed the molecular basis for RNA binding, substrate selection, dimerization, and the crucial role of RNA-binding domain 3 for ADAR1 editing.

Utilizing our well-established biochemical assays, we evaluated ZYS-1’s ADAR1 inhibitory activity using HT-dsRNA, derived from the physiologically relevant RNA substrate 5-HT_2C_R pre-mRNA ^14^, with direct inosine quantification. The compound showed no inhibitory activity at concentrations up to 1 mM (**Figure 1C-D**). To rule out the possibility that ADAR1 inhibition by ZYS-1 is dsRNA sequence-dependent, we also tested another dsRNA substrate, GLI-dsRNA-1 as used in *Nature Cancer* ^1^, but could not observe any appreciable inhibition of ADAR1 activity by ZYS-1 up to 1 mM (**Figure 1C-D**).

Collectively, these results demonstrate that the ADA assay kit used by Wang et al. does not reflect ADAR1-specific activity, and ZYS-1 is unable to inhibit ADAR1 in well-validated biochemical assays.

While the DARTS and CETSA data suggest ZYS-1 interacts with ADAR1, these assays can only reflect binding with ADAR1 but cannot confirm direct enzymatic inhibition. The lack of activity in our orthogonal ADAR1 biochemical assay, coupled with the reliance on an ADA-based assay, undermines the claim that ZYS-1 is a catalytic inhibitor. Additionally, the cellular effects of ZYS-1 (e.g., reduced GLI1 editing) could arise from off-target mechanisms.

Intriguingly, the authors demonstrated that ZYS-1 reduces ADAR1 protein levels in cancer cells through a post-transcriptional degradation mechanism independent of proteasome, autophagy/lysosome pathways, and cell death. It is possible that ZYS-1 exerts its effects by affecting protein translation. We are curious about the proteome changes upon ZYS-1 treatment in cells.

The SPR-derived equilibrium dissociation constant (*K*_d_) of 0.313 μM for ZYS-1 binding to ADAR1 (Fig. 5e) raises additional questions. The authors fit maximal response values to a concentration series to calculate *K*_d_, rather than using kinetic parameters (*k*_on_ and *k*_off_). Without sensorgram data, it is unclear whether binding reached equilibrium or if the observed responses reflect specific interactions. Again, the SPR experiment only reflects binding instead of inhibition of ADAR1.

In light of these concerns, we urge the authors to provide further validation using ADAR1-specific biochemical assays (e.g., RNA editing assays) and to disclose raw SPR sensorgrams and experimental details. Clarifying these points is essential to establish ZYS-1 as a genuine ADAR1 inhibitor and to advance its potential therapeutic applications.

## Supplementary Information

### Materials

HT-dsRNA, GLI-dsRNA-1, and GLI-dsRNA-2 were synthesized using an in vitro transcription method, and the RNAs were purified by 15% denaturing (7 M urea) polyacrylamide gel electrophoresis (PAGE), extracted, precipitated by ethanol, air-dried, and dissolved with RNase-free H_2_O. GLI-dsRNA-Product was obtained from the Keck Biotechnology Resource Laboratory at Yale University and purified using cartridge purification.

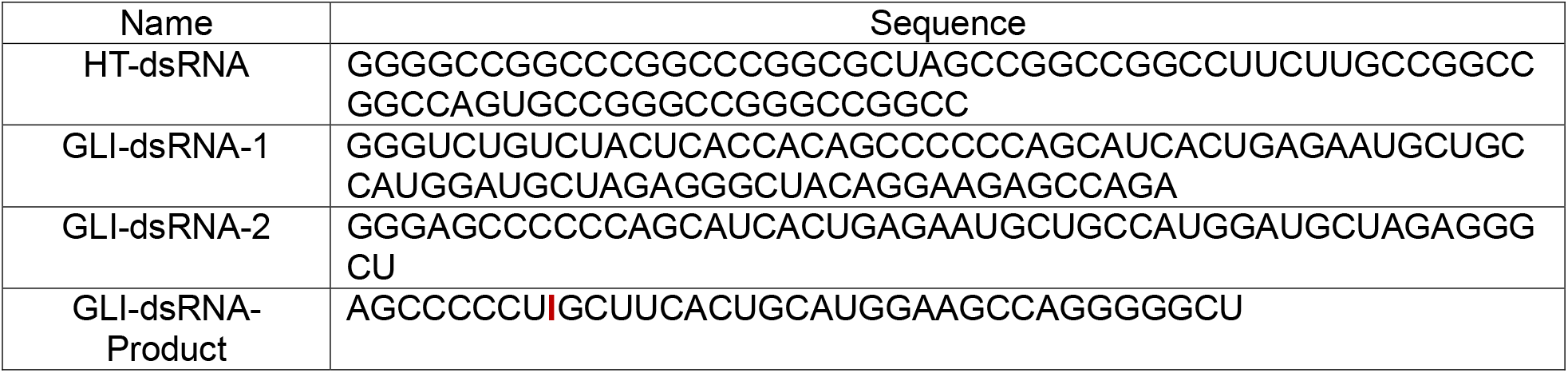

The gene of human ADAR1 was obtained from Addgene (plasmid #117927). The ADAR1 expression plasmid was constructed using the In-Fusion Snap Assembly method (Takara), with the ADAR1 gene inserted into a modified pLEXm mammalian expression vector containing an N-terminal MBP tag. ADAR1 was expressed in HEK293T suspension cells (ATCC, catalog CRL-3216). Transfection was carried out using PEI (Polysciences, catalog 23966-1) along with plasmid DNA. The protein was purified using amylose resin, followed by size-exclusion chromatography with Superose S6. Detailed methods can be found in Deng et al., Biochemical Profiling and Structural Basis of ADAR1-Mediated RNA Editing ^13^.

ZYS-1 was purchased from ChemScene and further verified in-house by ^1^H NMR (600 MHz, DMSO-*d*_6_) δ 8.34 (s, 1H), 7.89 (d, *J* = 44.2 Hz, 2H), 5.84 (d, *J* = 5.6 Hz, 1H), 5.62 (d, *J* = 6.0 Hz, 1H), 5.48 (d, *J* = 5.2 Hz, 1H), 4.67 (q, *J* = 5.6 Hz, 1H), 4.14 (dq, *J* = 58.7, 4.8 Hz, 2H), 3.94 (dd, *J* = 11.6, 5.1 Hz, 1H), 3.84 (dd, *J* = 11.6, 6.3 Hz, 1H).

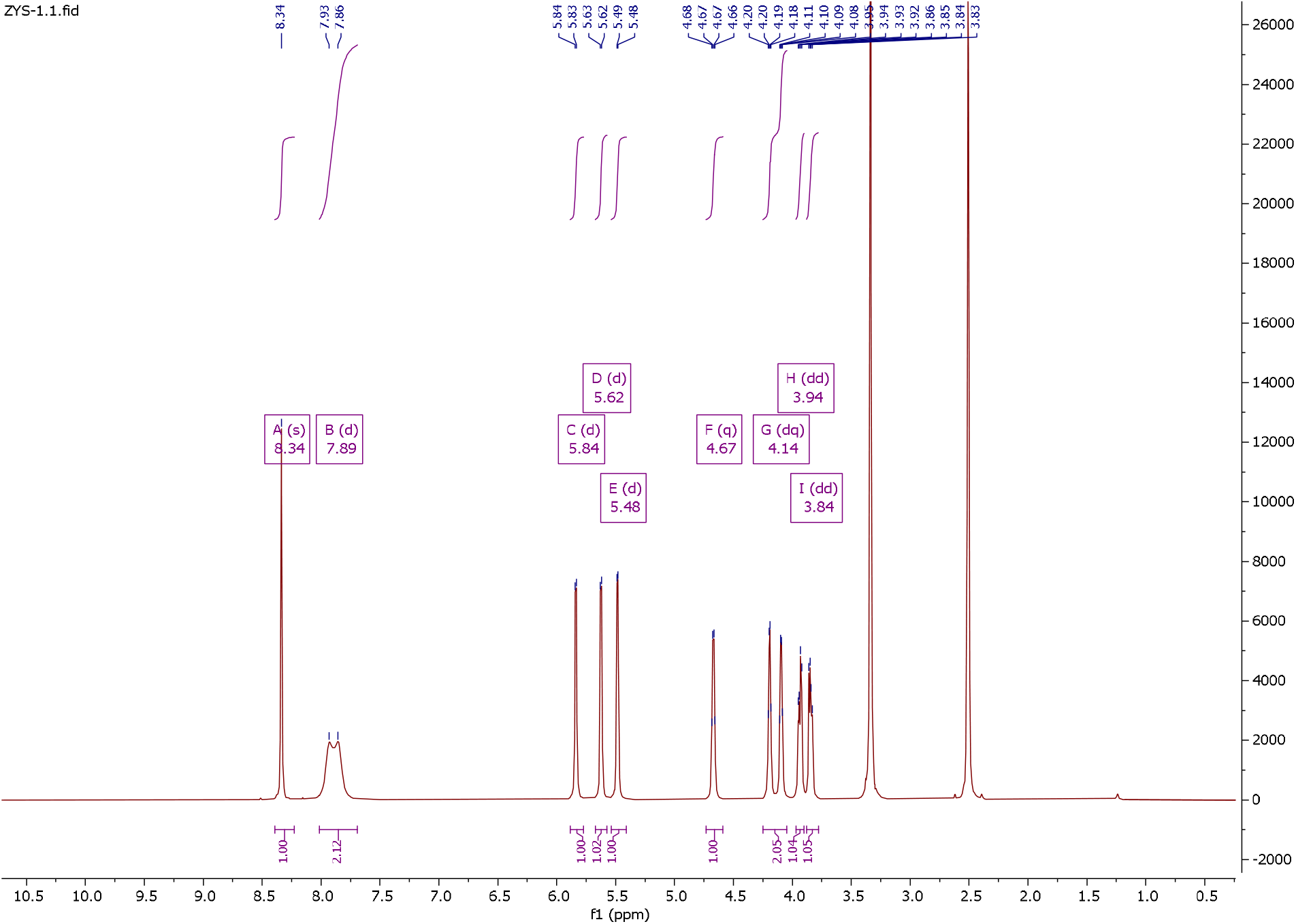

Adenosine Deaminase Activity Assay kit (Fluorometric, Abcam, ab204695)

### Adenosine Deaminase Activity Assay

To evaluate the assay response, we prepared standard curves using both free inosine and inosine-containing dsRNA substrates. Briefly, inosine standard (10 μM) or synthetic inosine-containing dsRNA GLI-dsRNA-Product (10 μM, GLI-dsRNA-2) was serially diluted in ADA Assay Buffer to achieve final concentrations of 0, 0.4, 0.8, 1.2, 1.6, and 2.0 μM in a total volume of 50 μL per well in a 96-well plate.

The standards were transferred to a flat-bottom white 96-well plate (50 μL per well). To each well, we added 50 μL of Background Control Mix containing 45 μL ADA Buffer, 2 μL ADA Converter, 2 μL Converter Enzyme VIII, and 1 μL OxiRed Probe. After thorough mixing, fluorescence was measured kinetically (Ex/Em = 535/587 nm) at 37°C for 60 minutes using a CLARIOstar plate reader. We analyzed the fluorescence values within the linear response range (23-33 minutes post-reaction initiation), recording the relative fluorescence units (RFU) at two timepoints (RFUS1 and RFUS2). Background readings from the 0 μM standard wells were subtracted from all measurements. The resulting values were plotted against standard concentrations, and linear regression analysis was performed using GraphPad software to establish standard curves for both free inosine and inosine-containing dsRNA.

### ADAR1 Editing Activity Assay

The in vitro RNA editing assay was performed by incubating 150 nM ADAR1 with 50 nM of HT-dsRNA (derived from 5-HT_2C_R pre-mRNA) or GLI-dsRNA (derived from Glioma-associated oncogene 1 mRNA) substrate in a reaction buffer containing 10 mM HEPES-K (pH 7.5), 70 mM KCl, 5% glycerol, 1 mM DTT, and the indicated concentration of the ZYS-1 compound at 37°C for 30 minutes. Please note the catalytic efficiency of ADAR1 is very low. In our assay condition, it is essentially a single turnover enzyme. Following the editing reaction, RNA was purified using the RNA Cleanup Kit (NEB, T2030L). The purified RNA was then incubated with 100 nM EcEndoV and 2 mM MnCl_ at 37°C for an additional 30 minutes. The reaction was terminated by adding 2× loading buffer (93.5% formamide, 0.025% xylene cyanol FF, and 50 mM EDTA, pH 8.0) and heating at 85°C for 5 minutes. Quenched reactions were resolved on 15% denaturing polyacrylamide gels (Urea-PAGE). The gels were then exposed to a phosphorimager plate for 1 hour and imaged using the Sapphire Biomolecular Imager (Azure Biosystems). Assays were performed in three independent replicates, and the band intensities of substrates and products were analyzed using Azure Spot (Azure Biosystems) and plotted using GraphPad Prism.

